# Differential adenine methylation analysis reveals increased variability in 6mA in the absence of methyl-directed mismatch repair

**DOI:** 10.1101/2022.12.14.520158

**Authors:** Carl J. Stone, Gwyneth F. Boyer, Megan G Behringer

## Abstract

Methylated DNA adenines (6mA) are an important epigenetic modification in bacteria that affect varied cell processes like replication, stress response, and pathogenesis. While much work has been done characterizing the influence of 6mA on specific loci, very few studies have examined the evolutionary dynamics of 6mA over long time scales. Utilizing third-generation sequencing technology, we produced a detailed analysis of 6mA methylation across the *Escherichia coli* K-12 substr. MG1655 genome. 6mA levels were consistently high across GATC sites; however, we identified regions where 6mA is decreased, particularly in intergenic regions, especially around the -35 promoter element, and within cryptic prophages and IS elements. We further examined 6mA in WT and methyl-directed mismatch repair-knockout (MMR-) populations after 2400 generations of experimental evolution. We find that, after evolution, MMR-populations acquire significantly more epimutations resulting in a genome-wide decrease in 6mA methylation. Here, clones from evolved MMR-populations display non-deterministic sets of epimutations, consistent with reduced selection on these modifications. Thus, we show that characterization of 6mA in bacterial populations is complementary to genetic sequencing and informative for molecular evolution.

## INTRODUCTION

DNA methylation introduces genetic variation in a conservative manner by altering the expression and function of DNA without permanently changing the sequence. Bacterial epigenetics, primarily in the form of DNA base methylation, has an outsized role in the regulation of various cellular processes^1–7^. The methylated DNA bases found in bacteria and archaea include 6mA, 5mC, and N^4^-methylcytosine (4mC), which are added by DNA methyltransferases either as part of a restriction-modification system or by solitary methyltransferases without associated restriction enzymes, called orphan methyltransferases^1, 4, 8–11^. In gammaproteobacteria, 6mA modified by the orphan methyltransferase Dam at GATC motifs is the most abundant modified base in the genome^12–15^. There are 19,120 of these palindromic GATC motifs in the *Escherichia coli* K-12 genome, or one GATC site on average every ∼250 bp^16^. 6mA influences various processes including, but not limited to, replication timing, transposon mobility, gene expression, virulence, and methyl-directed mismatch repair (MMR)^1, 6, 17–28^. Many phenotypes regulated by 6mA involve phase variation, such as the pathogenic fimbriae *pap* in uropathogenic *E. coli* and *std* in *Salmonella enterica* serovar Typhimurium^29–31^. In these phase-variable phenotypes, whether gene expression is on or off depends on the methylation status of multiple nearby regulator binding sites^32^. Thus, the short-term stability but long-term mutability of DNA methylation facilitates these phase-variable phenotypes by allowing for stable regulation of loci that can rapidly switch with epigenetic changes^33^.

The most widespread use of 6mA across the *E. coli* genome is in directing MMR. Methyl-directed MMR GATC sites are nearly all methylated on both strands, except for the brief period after chromosome replication before the daughter DNA strand is methylated by Dam^19, 34^. When a mismatch is detected by MMR proteins (MutHLS), the nearby hemimethylated GATC sites are used to direct the MMR protein complex to remove the mismatched base from the daughter strand^19, 23, 35, 36^. In E. coli, inactivation of any of the MutHLS proteins disrupts MMR and leads to a ∼150-fold increased mutation rate^37^. Since MMR relies on the presence of hemimethylated GATC sites, alterations to the function of Dam in *E. coli* and other gammaproteobacteria also reduce the effectiveness of MMR^18, 22, 38^.

Given the role of MMR in preventing replicative errors from becoming permanent mutations, we hypothesize that maintaining MMR imposes selective pressure on the *E. coli* genome to maintain GATC sites and associated 6mA modifications. While 6mA can have local gene expression effects when GATC sites coincide with regulatory elements and transcription factor binding sites, MMR is likely the most significant use of (and influence on) 6mA genome-wide. Thus, we predict that in the absence of MMR, *E. coli* experimentally evolved for thousands of generations will exhibit increased variability in per-site 6mA methylation and a genome-wide decrease in 6mA due to a sustained accumulation of epimutations that do not incur the strongly deleterious fitness effect of causing MMR dysfunction. There has been limited examination of the dynamics of 6mA in evolving bacterial populations^39, 40^. Thus, in this study, we sought to investigate how the presence and absence of MMR influence 6mA methylation in evolving bacterial populations over an extended time period. To accomplish this, we created a differential methylation-calling pipeline designed for bacterial 6mA data from third-generation sequencing: CoMMA (<u>C</u>omparison <u>o</u>f <u>M</u>icrobial <u>M</u>ethylated <u>A</u>denines). Third-generation sequencing platforms (such as PacBio’s SMRT sequencing or ONT’s Nanopore sequencing) allow for accurate and cost-effective identification of DNA base modifications. Using Nanopore sequencing and CoMMA, we first characterized genome-wide methylation frequency at GATC sites in *E. coli* MG1655. Then, focusing on clones isolated from evolved populations with initially WT and MMR-deficient genetic backgrounds, we characterized how 6mA methylation frequency changed among all 19,120 GATC motifs after 2000 generations of experimental evolution. We then compared our findings to studies of spontaneous mutation rates in *E. coli* to determine how MMR constrains genetic and epigenetic evolution at GATC sites. Together, our results illustrate the evolvability of 6mA and its potential to contribute to heritable change in bacterial populations.

## RESULTS

### 6mA methylation status varies between genomic features in *E. coli* MG1655

We first measured genome-wide methylation in *E. coli* K-12 MG1655 (strain PMF2) to characterize methylation trends in our wild-type ancestor. Briefly, three colonies from an overnight culture were sequenced on an ONT MinION portable sequencer. Nanopore sequencing produced an average read depth of ∼50 reads per GATC site, and there was no significant strand bias in coverage (**Fig. S2**). To measure methylation from Nanopore reads, we used the methylation classification program Megalodon to calculate the proportion of methylated reads to total reads at each GATC site, called the percent methylation (**Dataset S1**)^41^. Previous benchmarking studies have shown Megalodon to be one of the most reliable methylation callers when applied to data generated from Nanopore sequencing^41–43^. To confirm this we compared the performance of multiple independently developed methylation calling programs, and with our data Megalodon consistently identified most of the same sites as methylated or unmethylated and correctly identified stably unmethylated sites reported previously (see **Methods, Supplementary Text 1, Fig. S3, Fig. S4, Table S1, Table S2**)^34, 44–47^. After classifying methylated sites with Megalodon, all GATC sites were annotated with data from RegulonDB, a curated database of *E. coli* K-12 transcriptional regulatory networks^48^. Consistent with previous reports, most GATC sites were almost completely methylated, with a median genome-wide percent methylation of 97% (**Fig. 1A**)^8, 34, 39, 49^. Methylation profiling revealed that 177 double-stranded GATC sites are hypomethylated on at least one strand, of which 138 were hemimethylated and 39 were hypomethylated on both strands (**Table S3**). Intergenic GATC sites were significantly less methylated compared to those within coding sequences (median percent methylated reads of 93% compared to 97%; Wilcoxon rank sum test, P < 2.2×10^-16^). This decrease at intergenic sites is likely due to the higher proportion of transcriptional regulatory regions and transcription factor binding sites in intergenic regions, where these sites may exclude binding of the Dam methyltransferase to GATC sites.

**Figure 1.**
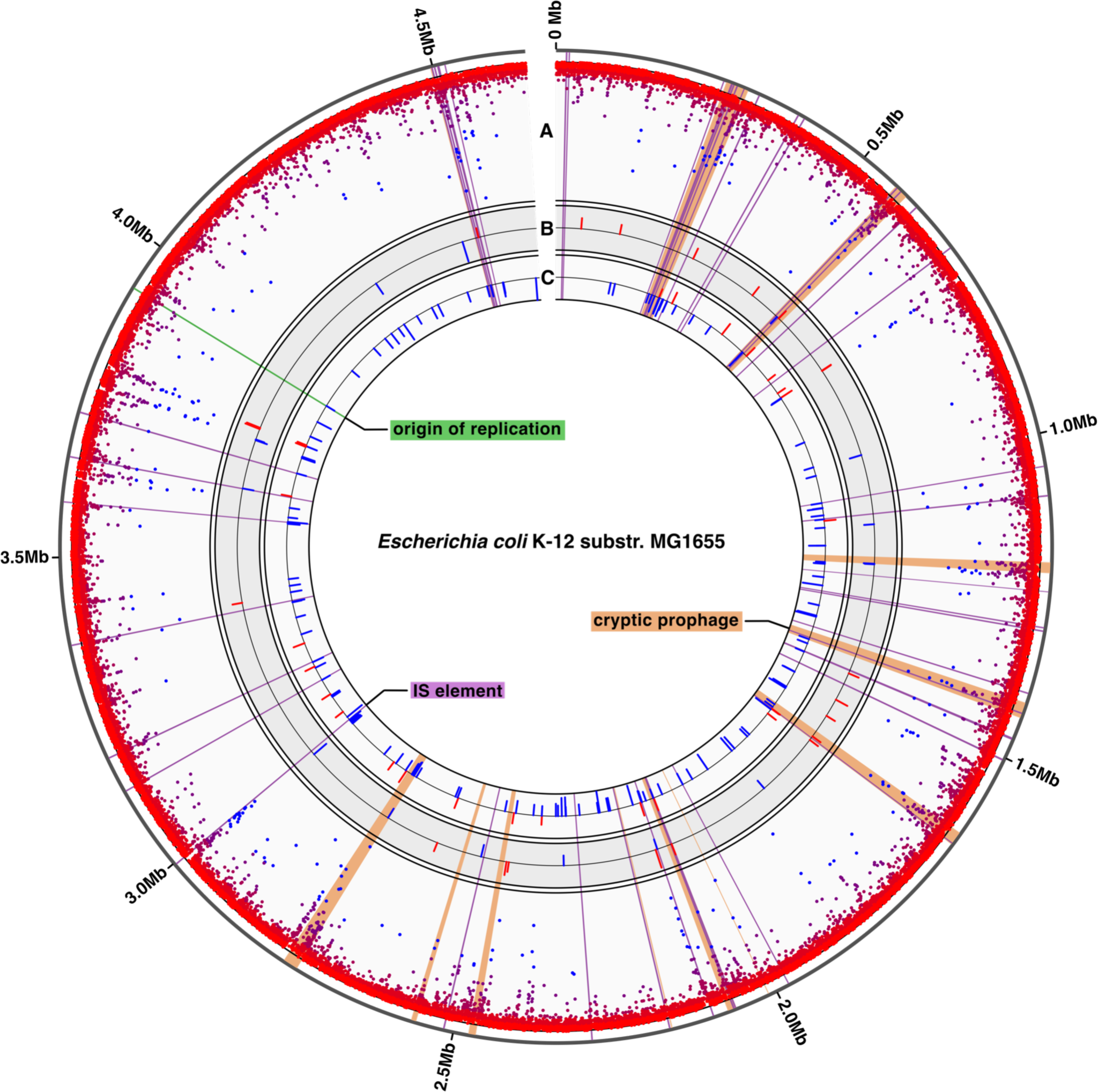
Genome-wide 6mA methylation in *E. coli* K-12 substr. MG1655. The origin of replication (green), IS elements (purple) and cryptic prophages (orange) are shown for reference. (A) Percent methylation at GATC sites. Each point represents one GATC site and points are colored to reflect percent methylation from 0% (blue, inside) to 100% (red, outside). (B-C) Differentially methylated GATC sites in evolved WT (A) and MMR-(B) clones. Red bars are sites with increased methylation and blue bars are decreased methylation, and the height of the bar reflects the difference in percent methylation between the ancestor and the evolved clones. Each ring ranges from -25% to 25%.

For a broader view of the factors that influence GATC methylation and to determine if hypomethylation was associated with GATC sites that may be shielded by protein binding, we looked at sites within intergenic regions where DNA-protein interactions likely occur, such as sigma factor binding sites and binding sites of other transcriptional regulatory proteins^2, 33, 46, 47, 50^. Overall, within these intergenic regions, there is a general decrease in methylation around promoters. GATC sites within -10 and -35 promoter elements were significantly less methylated (95.2% for -10 elements, P_-10_ = 6.7×10^-8^; 90.4% for -35 elements, P_-35_ = 6.7×10^-10^, Wilcoxon signed rank test with Benjamini-Hochberg correction) than the genome-wide median percent methylation (96.6%). This is best observed when methylation is averaged across all promoter regions in the genome; there is a lower frequency of methylated reads around 100 bp before and after the transcription start site, with a maximum decrease in methylation of ∼5% centered on the -35 element (**Fig 2A**). Moreover, -35 promoter elements known to be bound by σ^70^, σ^38^, and σ^24^ had significantly lower percent methylation than the genome-wide median (Wilcoxon signed rank test with BH correction: P_sigma70_ = 2.0×10^-10^; P_sigma38_ = 7.0×10^-4^; P_sigma24_ = 1.6×10^-5^) (**Fig 2B**).

**Figure 2.**
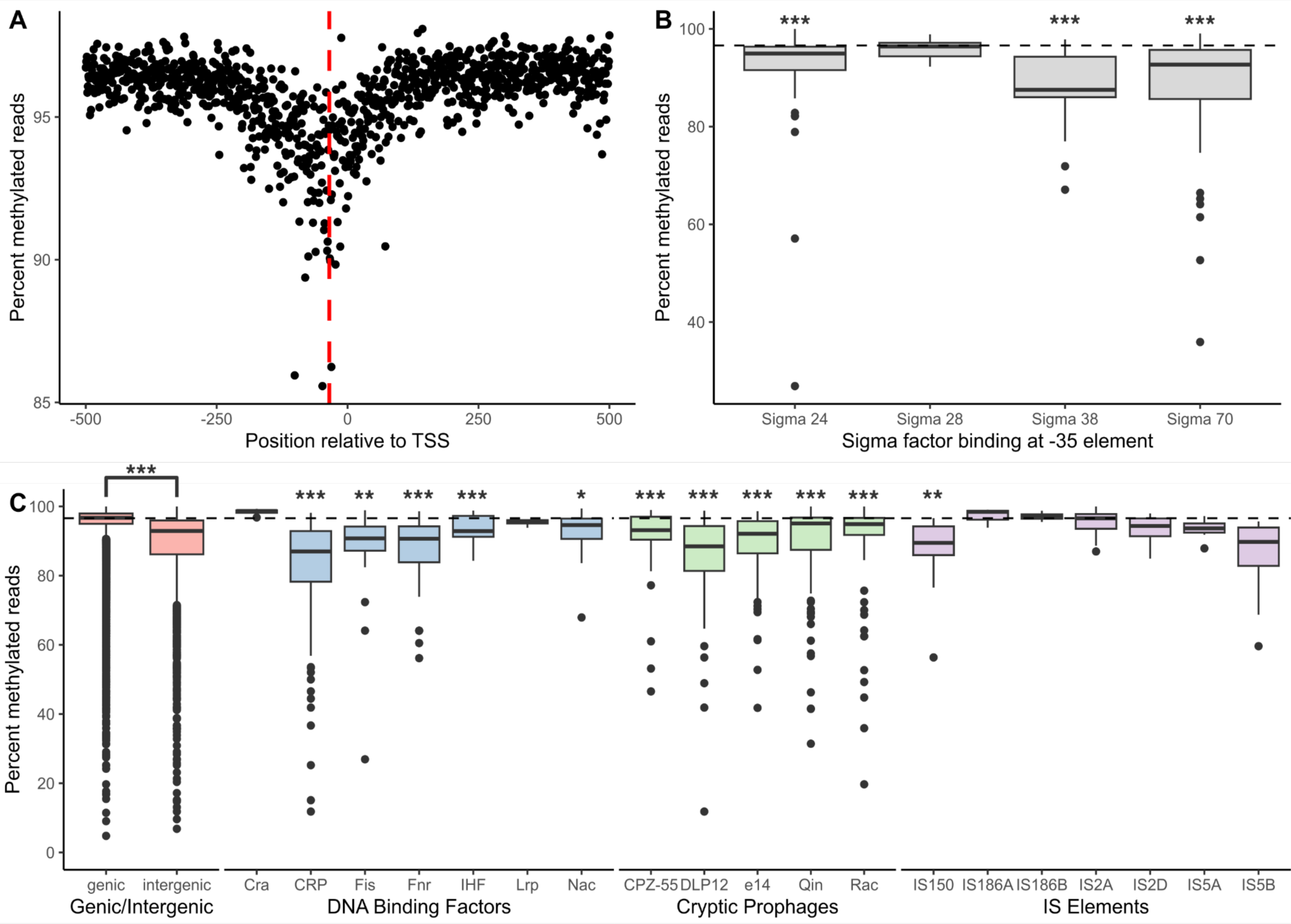
Genome-averaged methylation decreases around core promoter elements. (A) Median methylation at GATC sites aligned relative to the transcription start sites (TSS) for all promoters. Points represent the median methylation for all GATC sites at that position. The -35 position relative to the TSS is shown for reference (dashed red line). (B) Comparison of percent methylated reads at -35 elements by specific sigma factor binding. Boxplots show the median, first quartile, and third quartile. Sigma factors were compared to the genome-wide median percent methylation (97%, dashed line) with a one-sample Wilcoxon signed rank test with Benjamini-Hochberg correction. P-values are shown in B and C as *** < 0.001 < ** < 0.01 < * < 0.05. (C) Different genomic features showed variable methylation status. Percent methylation of GATC sites within each feature type was compared to the genome-wide median percent methylation (97%, dashed line) with one-sample Wilcoxon signed rank tests with Benjamini-Hochberg correction.

In addition to core promoter elements, binding sites of other transcriptional regulatory proteins within intergenic regions also exhibited differences in their degree of hypomethylation (**Fig 2C**). Specifically, the global regulators CRP, Fis, Fnr, and Nac were significantly less methylated than the genome-wide median (Wilcoxon signed rank test with BH correction: P_CRP_ = 2.0×10^-18^; P_Fis_ = 4.2×10^-3^; P_Fnr_ = 9.1×10^-6^; P_Nac_ = 4.98×10^-2^). Differences in methylation were also observed within coding regions for a few genomic features (**Fig 2C**). Across the 12 cryptic prophages in the *E. coli* K-12 genome, significantly decreased methylation is observed in 9 of them: CP4-44, CP4-57, CP4-6, CPS-53, DLP12, e14, Qin, Rac, and PR-Y (CP4-44: 94.0% methylation, P = 6.2×10^-4^; CP4-57: 92.3%, P = 9.4×10^-16^; CP4-6: 95.3%, P = 2.5×10^-13^; CPS-53: 92.4%, P = 4.7×10^-6^; DLP12: 88.5%, P = 3.7×10^-13^; e14: 92.1%, P = 1.1×10^-8^; Qin: 95.1%, P = 2.9×10^-6^; Rac: 94.9%, P = 4.3×10^-8^; PR-Y: 93.8%, P = 2.6×10^-2^; Wilcoxon signed rank test with BH correction)^51, 52^. Of the various IS elements, *IS*150, *IS*4, *IS*30A, *IS*30C, and *IS*30D had significantly decreased methylation (*IS*150: 89.4% methylation, P = 2.6×10^-3^; *IS*4: 94.1%, P = 1.3×10^-2^; *IS*30A: 89.7%, P = 4.3×10^-2^; *IS*30C: 85.8%, P = 1.5×10^-2^; *IS*30D: 86.7%, P = 2.2×10^-2^)^53–55^. Lastly, some genomic features, such as REP elements and terminator sequences, do not show measurable differences in 6mA methylation compared to the genome-wide average. However, these regions either do not contain many GATC sites or are poorly annotated. Comparing sequencing replicates from future studies across a variety of conditions, combined with improving genomic annotations, could allow for the detection of higher-resolution methylation differences and the characterization of methylation on abundant but unannotated sites.

### Differential methylation analysis shows that MMR-clones evolve decreased GATC methylation

After characterizing a baseline *E. coli* K-12 substr. MG1655 6mA methylome, we compared GATC methylation across the experimental ancestor PMF2 strain and eight isolated clones from populations that were experimentally evolved for 2400 generations with either intact (WT) or disrupted mismatch repair (MMR-)^56^. Briefly, CoMMA adapts the differential methylation package methylKit to identify GATC sites with significant differences in methylation status between bacterial samples using logistic regression^57^. For a broad view of how methylation evolves, we focused on two populations from each of the WT or MMR-genetic backgrounds; from each of these populations, we selected two clones and ensured each clone belonged to a different evolved subpopulation. This sampling scheme would allow us to evaluate how methylation evolves within and among populations depending on their ability to conduct methyl-directed mismatch repair.

As with the ancestor, experimentally evolved clones were Nanopore sequenced, and methylated sites were called using Megalodon (**Dataset S2**). The median genome-wide methylation percentage of both genetic backgrounds decreased during experimental evolution with MMR-lines exhibiting a significantly larger median decrease in methylation (-0.15%) compared to evolved WT lines (-0.03%; Wilcoxon rank sum test, P < 2.2×10^-16^, **Fig. S5**). Next, we determined which specific GATC sites were differentially methylated between the evolved lines and the ancestor (**Fig 3A-B**). Using cutoffs P < 0.05 and percent methylation change >10%, there were significantly more differentially methylated sites in the MMR-lines (28 sites with increased methylation, 141 sites with decreased methylation) compared to the evolved WT lines (22 increased, 16 decreased; Fisher’s exact test, P < 2.2×10^-16^).

**Figure 3.**
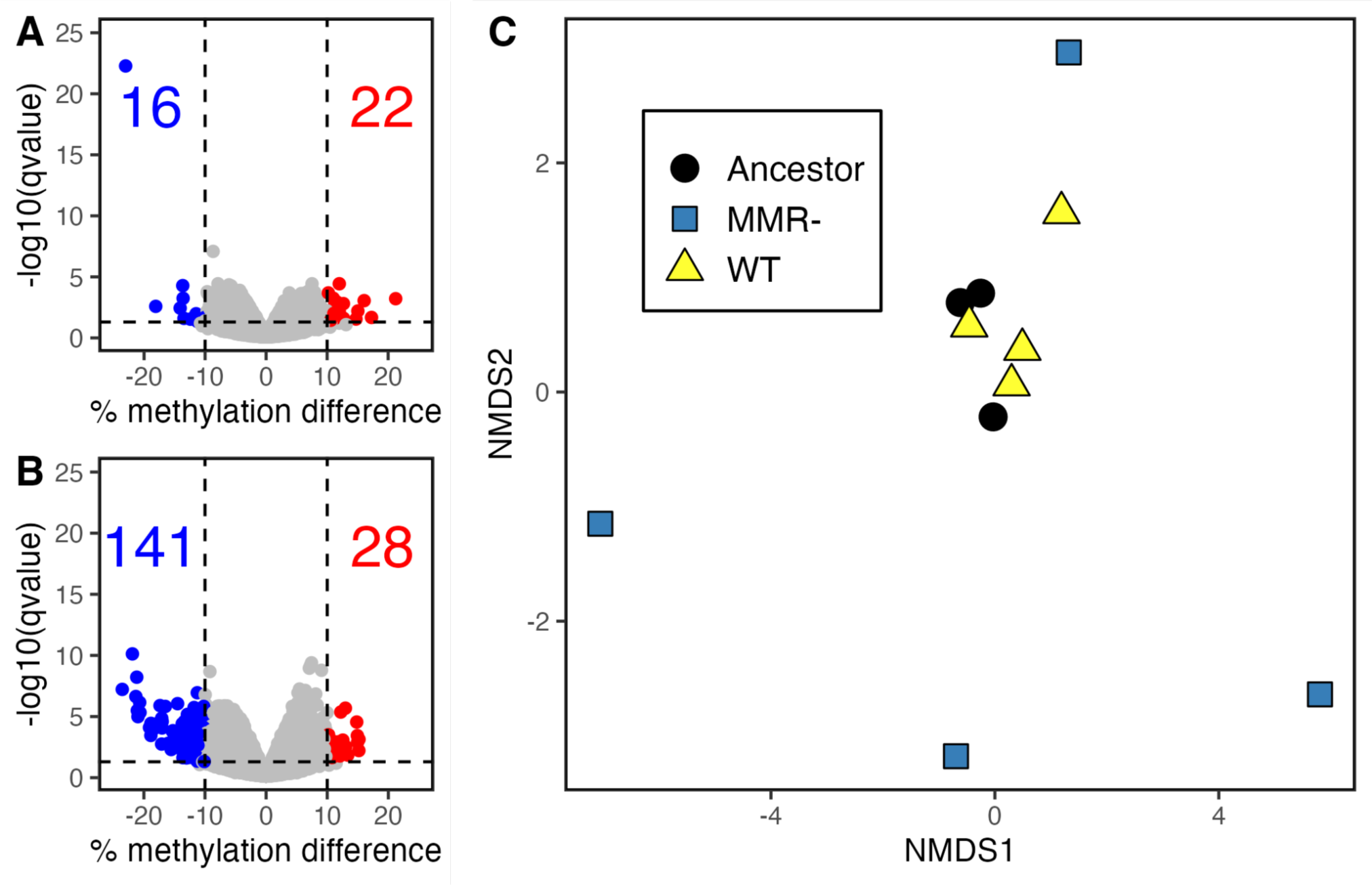
Differentially methylated GATC sites between the ancestor and experimentally evolved (A) WT clones or (B) MMR-clones. Differential methylation was called for WT and MMR-clones compared to the ancestor strain using methylKit. Differential methylation was determined for each site as a percent methylation difference compared to the ancestor greater than 10% and a q-value less than 0.05. Numbers shown in each plot field represent the number of differentially methylated sites that increased (red) or decreased (blue) in methylation in the evolved condition compared to the ancestor. (C) Comparison of methylation across all sites between the ancestor and experimentally evolved clones. Euclidean distance between clones was calculated using normalized β-values and ordination was performed with NMDS. Each point represents one clone.

Both evolved backgrounds also had a significant overrepresentation of differentially methylated sites within certain genomic regions, especially in sites where DNA-protein interactions occur. Across the genome, differentially methylated sites in both WT and MMR-evolved clones appear to coincide with regions of low methylation in the ancestor and with cryptic prophages and IS elements (**Fig. 1B-C**). Sites that were hemimethylated and hypomethylated in the MG1655 ancestor were enriched for differential methylation in the evolved clones (Fisher’s exact test, P < 2.2×10^-16^; **Table S3**). Differentially methylated sites in both evolved backgrounds were significantly over-enriched in transcription factor binding motifs and IS elements (Fisher’s exact test with FDR correction, P < 0.05, **Table 1**). Sites with increased methylation in both evolved backgrounds were also over-enriched within cryptic prophages, and sites with decreased methylation were over-enriched in Rho-independent terminators. The MMR-evolved lines also showed enrichment for under-methylated sites within cryptic prophages (P < 2.2×10^-16^) and within promoters, specifically σ^70^-binding sites in the -35 region (P < 2.2×10^-16^).

**Table 1.** Overrepresentation of differentially methylated adenines in protein-DNA interaction motifs.

| Feature Type | # GATC sites<br>in ancestor | Evolved WT |  |  |  | Evolved MMR- |  |  |  |
| --- | --- | --- | --- | --- | --- | --- | --- | --- | --- |
|  |  | Increased<br>methylation |  | Decreased<br>methylation |  | Increased<br>methylation |  | Decreased<br>methylation |  |
|  |  | n | p-value | n | p-value | n | p-value | n | p-value |
| Transcriptional regulator binding sites | 313 | 7 | 0.095 | 12* | < 0.001 | 11^ | 0.014 | 58* | < 0.001 |
| Origin of replication | 22 | 0 | 1.000 | 0 | 1.000 | 0 | 1.000 | 1 | 0.647 |
| -10 element | 116 | 2 | 0.465 | 2 | 0.428 | 4 | 0.187 | 6 | 0.299 |
| -35 element | 60 | 1 | 0.590 | 1 | 0.572 | 1 | 0.694 | 12* | < 0.001 |
| Sigma 24 binding sites | 67 | 2 | 0.294 | 1 | 0.590 | 2 | 0.380 | 4 | 0.299 |
| Sigma 28 binding sites | 11 | 0 | 1.000 | 0 | 1.000 | 0 | 1.000 | 0 | 1.000 |
| Sigma 32 binding sites | 2 | 0 | 1.000 | 0 | 1.000 | 0 | 1.000 | 0 | 1.000 |
| Sigma 38 binding sites | 17 | 0 | 1.000 | 0 | 1.000 | 0 | 1.000 | 3* | 0.043 |
| Sigma 70 binding sites | 86 | 1 | 0.694 | 2 | 0.313 | 3 | 0.254 | 12* | < 0.001 |
| Genes | 35595 | 312 | 1.000 | 275 | 1.000 | 442 | 1.000 | 952 | 1.000 |
| Rho-dependent terminators | 8 | 0 | 1.000 | 0 | 1.000 | 0 | 1.000 | 1 | 0.380 |
| Rho-independent terminators | 46 | 1 | 0.514 | 4* | 0.003 | 0 | 1.000 | 6* | 0.011 |
| IS elements | 283 | 10^ | 0.002 | 10* | 0.001 | 11^ | 0.007 | 18* | 0.014 |
| Cryptic prophages | 795 | 15^ | 0.038 | 11 | 0.249 | 24^ | 0.001 | 62* | < 0.001 |
| No Feature | 1191 | 35^ | < 0.001 | 30* | < 0.001 | 44^ | < 0.001 | 134* | < 0.001 |

P-values were calculated by one-tailed Fisher's exact test followed by BH correction. ^ = significant number of sites with increased methylation. \* = significant number of sites with decreased methylation.

Gene ontology (GO) enrichment analysis of genes containing or near differentially methylated sites revealed enrichment in genes associated with stress response, biofilm formation, and virulence factor production. Of the 194 differentially methylated sites identified between both WT and MMR-evolved clones, 13 sites were in the same gene or intergenic region between the two genotypes (**Fig. 4A**). Ten of these sites were in the same palindromic GATC site, and 7 of those were in the same adenine residue. GO enrichment analysis was then done on genes that contained differentially methylated GATC sites, or if the GATC site was not within a gene then the nearest gene was used. Genes that were less methylated in the MMR-clones were significantly enriched for biofilm formation and transcriptional regulators (**Fig. 4B**). Genes that were more methylated after evolution were enriched for biofilm formation in the evolved WT clones and for lipopolysaccharide metabolism in both WT and MMR-clones. Interestingly, six of the fifteen biofilm genes identified belong to three different putative chaperone-usher fimbrial operons. These fimbriae are cryptic in *E. coli* K-12 under laboratory conditions, but when overexpressed they increase biofilm formation^58^. Other biofilm genes *yfaL* and *ycgV* are homologs of the cryptic prophage-encoded autotransporter and adhesin Antigen 43 (Ag43 encoded by *flu*)^59^. Both fimbriae and Ag43 are known to be subject to methylation-regulated phase variation in *E. coli* and related species^31, 59–61^. Other enriched gene ontologies include lipopolysaccharide synthesis (including the non-functional O-antigen polymerase *wbbH*), acid stress response genes, phage defense genes, and various transcriptional regulators involved in stress response (**Table S4**)^62–65^.

**Figure 4.**
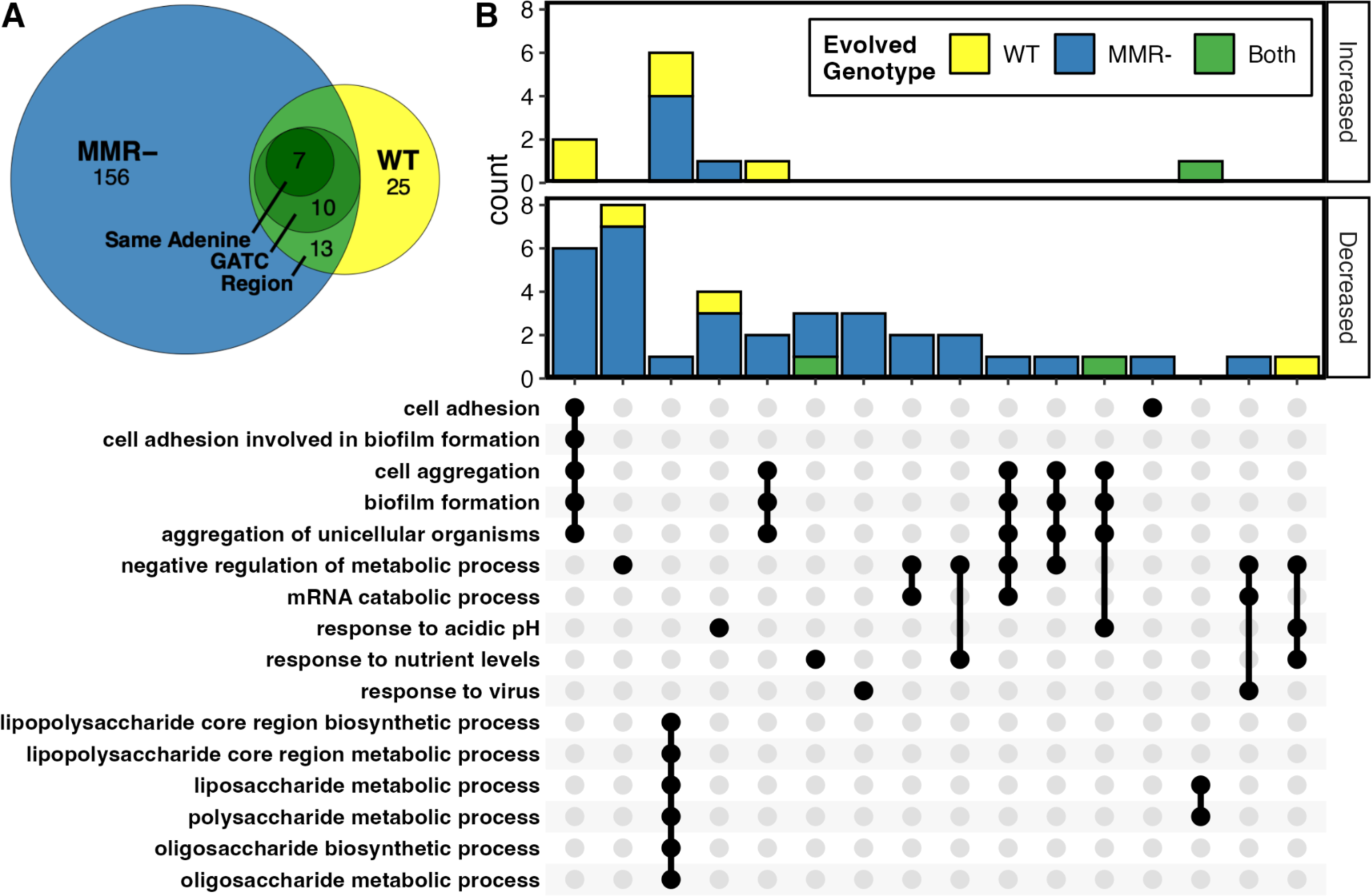
Gene ontology enrichment analysis of differentially methylated GATC sites. (A) Intersect between differentially methylated sites in evolved WT and MMR-genotypes. Sites in the same region are located in the same gene or the same intergenic region. (B) UpSet plot of significantly enriched gene ontology terms of genes associated with differentially methylated GATC sites that increased or decreased in methylation in evolved WT, MMR-, or both. Bar heights are the number of genes associated with the intersecting GO terms shown below.

Comparison of the full 6mA methylomes between evolved clones and their ancestor revealed a non-deterministic change in 6mA in the absence of MMR consistent with random genetic drift. The percent of methylated reads at all GATC sites was compared between samples by clustering using non-metric multidimensional scaling (NMDS; **Fig. 3C**). The evolved WT clones and their ancestor clustered together, showing that their 6mA methylomes remain similar after extended evolution (Pairwise PERMANOVA Ancestor vs. WT, P = 0.168; Analysis of multivariate homogeneity of group dispersions, ANOVA with Tukey’s HSD: P_Ancestor-WT_ = 0.99). In contrast, MMR-evolved clones diverge from the ancestor in random directions, and the MMR-clones do not cluster together. The centroid of the MMR-clones was near that of the ancestor and WT clones but significantly different than the WT clones, and MMR-clone dispersions were significantly greater than both the ancestor and WT groups (Pairwise PERMANOVA WT vs. MMR-, P = 0.026; PERMDISP2, ANOVA with Tukey’s HSD: P_Ancestor-MMR-_ = 0.018, P_WT-MMR-_ = 0.010). This suggests that 6mA differences that occur during evolution in MMR-lines are non-deterministic, resulting in significant changes to methylation for each sample but not strongly in any one direction for the whole group. MMR-clones from the same population did not cluster together, indicating that shared evolutionary history does not have a large effect on broad 6mA differences over this time scale. Together, the significant number of under-methylated sites and the random divergence of differentially methylated sites in the MMR-clones compared to the ancestor suggest that 6mA is shaped by genetic drift in the absence of MMR.

### MMR-evolved lines accumulate GATC mutations at a rate that indicates relaxed selection

After identifying decreased maintenance of 6mA in the absence of MMR, we sought to genomically identify whether GATC sites in MMR-clones were also under decreased evolutionary constraint. Here, we characterized the fraction of mutations that altered GATC sites across the experimentally evolved clones from both evolved genetic backgrounds. After ∼2400 generations, MMR-clones had acquired a total of 959 mutations with 34 mutations (3.5% of mutations) causing a gain or loss of a GATC site (**Table 2, Table S4**). In contrast, the WT clones accumulated a total of 33 mutations, none of which had affected GATC sites. Since the relative lack of mutations in WT clones limited our statistical power, we compared the mutation rates of the experimentally evolved clones to mutation rates calculated in a minimally-selective mutation accumulation (MA) study by Lee *et al.*^37^. As this study directly measured the mutation rates of the same WT and MMR-PFM2 strain, we are able to accurately determine the expected rate of GATC mutation in the absence of selection^37^.

**Table 2.** Number of mutations in GATC sites in evolved clones.

| Background | Population* | Clones^ | Mutations | GATC<br>Gain/Loss | Percent GATC Mutated<br>(This Study) | Percent GATC Mutated<br>(Lee et al, 2012) |
| --- | --- | --- | --- | --- | --- | --- |
| MMR | 113 |  | 257 | 13 | 0.051 |  |
|  |  | 113-1 | 121 | 3 | 0.025 |  |
|  |  | 113-2 | 96 | 3 | 0.031 |  |
|  | 126 |  | 46 | 1 | 0.022 |  |
|  |  | 126-2 | 313 | 11 | 0.035 |  |
|  |  | 126-4 | 126 | 3 | 0.024 |  |
|  | Total |  | 959 | 34 | 0.035 (0.028 - 0.040) | 0.035 (0.028 - 0.043) |
| WT | 125 |  | 4 | 0 | 0 |  |
|  |  | 125-6 | 1 | 0 | 0 |  |
|  |  | 125-7 | 7 | 0 | 0 |  |
|  | 129 |  | 4 | 0 | 0 |  |
|  |  | 129-1 | 8 | 0 | 0 |  |
|  |  | 129-3 | 9 | 0 | 0 |  |
|  | Total |  | 33 | 0 | 0 (0.0 - 0.0) | 0.043 (0.025 - 0.067) |

\*Mutations shared between clones and occurred before subpopulation divergence
^Mutations unique to clones and occurred after subpopulation divergence

Reanalysis of the MA data to calculate mutation rates in tetranucleotide contexts determined that 3.5% of mutations that arose in MMR-lines during MA were in GATC sites, and is equal to the fraction of mutations affecting GATC sites in the MMR-lines during experimental evolution^37^. The similarity of these GATC mutation rates suggests that experimentally evolved MMR-clones acquire mutations in GATC sites at the neutral rate and may be evolving under relaxed selection. Additional evidence that GATC sites and associated adenine methylation may be evolving under relaxed selection in experimentally evolved MMR-clones include that the clones isolated from the MMR-Population 113 both acquired mutations resulting in a nonsynonymous D20A substitution proximal to the GATC recognition domain of *dam* methyltransferase^66^. While the exact consequences of this mutation have not been examined, this substitution could affect DNA binding and GATC recognition, indicating that perfect Dam function may not be necessary in the absence of MMR. Together, these data suggest that MMR is a major factor in maintaining GATC sites and *dam* methyltransferase, and in the absence of MMR these loci are vulnerable to mutation.

## DISCUSSION

In this study, we present what is to our knowledge a first-of-its-kind high-resolution characterization of the *E. coli* K-12 MG1655 6mA methylome that includes a functional and genic classification of 6mA methylation. We identified patterns of reduced methylation at intergenic regions, promoter elements, DNA binding sites, and cryptic prophages. This reduced methylation could result from the physical exclusion of Dam methyltransferase at these sites and potentially cause downstream effects on gene regulation and stress response. The ability to acquire high-resolution methylation data opens the door for future investigations on the specific effects of these methylation differences when paired with transcriptomic and biochemical experiments.

We also created and applied the CoMMA pipeline to investigate the effect of 2400 generations of evolution on WT and MMR-*E. coli* MG1655 clones. We found that the MMR-clones exhibited a significant decrease in methylation, indicating that 6mA is somewhat dispensable in the absence of MMR. While methylation was largely stable at the majority of GATC sites, MMR-clones exhibited significantly more differentially methylated sites than WT clones. Interestingly, these differentially methylated sites are often located in DNA binding sites. Adenine modification can affect specific binding affinity where 6mA coincides with a DNA binding site^67, 68^. Without the genome-wide pressure to maintain 6mA to minimize mutation rates, MMR-clones are free to alter 6mA at these sites with a bias toward decreased methylation, likely due to binding competition between transcription factors and the methyltransferase. Altered 6mA methylation that affects transcription factor binding can result in downstream effects and further alter cellular phenotypes. Our study demonstrates that differential methylomics is a powerful approach that can directly resolve differences between clones that are not apparent from genetic changes alone. Thus, the CoMMA pipeline is a useful tool that can complement genetic approaches.

An interesting observation from our study involves one of the MMR-evolved populations (P113), which acquired a mutation in the *dam* adenine methyltransferase resulting in a single amino acid substitution (DamD20A). The consequence of this mutation is unclear, although this residue is proximal to three S-adenosylmethionine-binding sites at K10, K14, and D54, and could potentially affect protein folding^66^. While there are no broad methylation differences within these clones compared to others of the same background, a DamD20A substitution may have contributed to the observed changes in methylation in these clones. Further studies are needed to investigate if this mutation altered Dam activity and how this affects genome-wide methylation patterns.

One potential impact of our findings is how 6mA methylation may influence the mutational spectrum and adaptive landscape of bacteria. DNA base modifications can affect the rates of spontaneous deamination; for instance, 5mC is vulnerable to deamination and contributes to the C -> T mutation rate^69–71^. Similarly, adenine and 6mA may deaminate to hypoxanthine, which pairs with cytosine during replication^72, 73^. While the spontaneous deamination rates of adenine versus 6mA have not been exhaustively studied, recent work identified a positive correlation between 6mA and A→G transitions in *Neisseria meningitidis*^74^. Thus, decreases in 6mA after MMR loss may result in a decreased A→G transition rate. This change to the mutational spectrum could potentially affect how MMR- or Dam-cells adapt to changing environments and may influence the trajectory of their evolutionary paths.

This study is the first-ever analysis of differential methylation between laboratory evolved bacterial lineages at this time scale. The ability to identify changes to 6mA at GATC sites over evolutionary time creates opportunities for the seamless integration of genetic and epigenetic data. Nanopore sequencing and pipelines like CoMMA are sufficiently reliable, affordable, and high-throughput to powerfully analyze genetic and epigenetic evolution in bacteria. As more studies uncover how bacterial DNA methylation affects gene expression, differential epigenetic analyses can be extended through time or between bacterial isolates and help to elucidate the relationship between mutation, epimutation, and gene expression as it relates to adaptation to challenging environments.^39, 40, 75, 76^.

## METHODS

### Bacterial strains and culture conditions

All bacterial strains were cultured and sequenced to identify mutations as part of a long-term experimental evolution study that has been described previously^56^. Briefly, the WT ancestor strain PFM2 is a prototrophic derivative of *E. coli* K-12 MG1655, and the MMR-deficient strain PFM5 is derived from PFM2 with an in-frame deletion of the *mutL* gene^56^. Long-term cultures were started by inoculating 10 mL of LB-Miller broth (BD Difco) in 16-× 100-mm glass culture tubes with a single colony of the progenitor strain^56^. Cultures were maintained shaking at 175 rpm, alternating transfers after 24 h of growth at 37 °C and 48 h of growth at 25 °C. Cultures of interest in this study were transferred by thoroughly vortexing the culture tube and inoculating 9 mL of LB-Miller broth with 1 mL of culture, resulting in a transfer of about 10^9^ cells^56^. Populations were grown and transferred for 2000 total generations of growth. After 2000 generations, samples from the evolving populations were streaked for isolation and eight random clones were saved in 40% glycerol at -80 °C for further and future experimentation.

### Nucleic acid isolation and Nanopore sequencing

Evolved and WT ancestor clones were revived from frozen storage on LB agar by incubating overnight at 37 °C. For each clone, a single isolated colony was randomly selected and restreaked on to fresh LB agar and again incubated overnight at 37 °C for DNA extraction. Following overnight incubation, colonies were collected from the agar using a 10mL loop and inoculated directly into DNA extraction buffer and DNA was further isolated with the UltraClean Microbial DNA Kit (Qiagen). Following recommendations from ONT, Extracted DNA was assessed for quantity using the Qubit High Sensitivity dsDNA kit and quality using the Genomic DNA ScreenTape on a TapeStation 4200 (Agilent). Libraries were then prepared for Nanopore sequencing using the PCR-free Rapid DNA sequencing Kit (SQK-RAD004, Nanopore) paired with the Rapid Barcoding Kit (SQK-RBK004). Prepared libraries were pooled and loaded on to a R.9.4.1 flow cell and sequenced on a MinION for 48h with live basecalling in MinKNOW deactivated.

### Bioinformatics and sequencing analysis

To determine the precision of methylation calling, we compared results for each of four methylation calling pipelines across three Nanopore sequencing replicates of the WT ancestor PFM2 and determined the number of inconsistently called sites. Initial Nanopore basecalling by converting fast5 files into fastq was conducted using GuppyGPU (v. 3.1.5+781ed57). Following basecalling, samples were demultiplexed with Guppy GPU. Average read length across all samples is 5583 +/- 275 nucleotides (+/- s.e.m). Following basecalling and demultiplexing, we evaluated four different pipelines (Tombo (v.1.5), mCaller (v.0.3), Deepsignal (v.0.1.6), and Megalodon (v.0.1.0)) for their precision and accuracy to identify base modifications and determine methylated adenines within GATC sites^41, 44, 77, 78^. Both Tombo and Megalodon were created by ONT for methylation calling, with Tombo being an older release that relies on statistical methods to annotate non-canonical bases and Megalodon being a newer release based on machine learning with deep neural networks^41, 78^. To increase accuracy, Megalodon can be run with a VCF file containing the position and identity of known variants, for which we used Illumina sequencing data for evolved clones that was previously published^55^. For pipelines not developed by ONT: DeepSignal applies a deep neural network to data that has been preprocessed by Tombo; and mCaller applies a deep neural network to data that has been preprocessed by Nanopolish. When running the mCaller pipeline we used Nanopolish v. 0.11.1). Across all samples, the mean read coverage across GATC sites was 57 reads per site (**Fig. S2**).

Identification of mutations affecting GATC sites in evolved clones was conducted using previously published WGS Illumina sequencing data. To identify the rate of mutations affecting GATC sites under minimal selection, we utilized previously published mutation accumulation data for the *E. coli* strains that served as the experimental ancestor for the WT (PFM2) and MMR- (PMF5, PMF2:Δ*mutL*)^37^. For both the experimentally evolved clones and the MA lines, mutations were called using GATK best practices and the genic location of mutations was annotated using SnpEff v. 4.3 by comparing VCF files of identified mutations for each clone to the *Escherichia coli* K-12 str. MG1655 reference genome (INSDC accession: U00096.3)^79^. To confirm a mutation occurred in a GATC site, the tetranucleotide context of each mutation was identified with the fill-fs -l 5 function from vcftools (v. 0.1.12) to annotate each variant with the flanking five upstream and downstream nucleotides^80^.

GATC sites were annotated as within different genomic features using all combined databases in RegulonDB v. 10.9^81^. These annotations were concatenated into a BED file, which was used to annotate GATC sites with the CoMMA function annotateMethylSites.

### Differential methylation calling

Differential methylation analysis was performed in R using CoMMA, a package that serves as a wrapper for methylKit v. 3.13^57^. GATC sites that had acquired mutations in any clone were removed from differential methylation analysis so that only GATC sites shared between samples were considered. To ensure the reliability of differential methylation analysis, CoMMA includes the following modifications to the standard methylKit workflow: 1) any sites covered by fewer than 10 reads were excluded from analysis; 2) methylation coverage at each site was normalized by median coverage; and 3) sites were ultimately considered to be differentially methylated based on a q-value cutoff of < 0.05 and percent methylation difference > 10%. Differential methylation was called separately between the ancestor and either the evolved MMR- or WT clones. Differential methylation was calculated by logistic regression with a sliding linear model (SLIM) to correct for multiple hypothesis testing^57, 82^.

### Statistical analysis and ordination

Statistical analysis comparing percent methylation (or ꞵ-value) was performed on M-values to remove the heteroscedasticity observed in ꞵ-value distributions^83^. M-values were calculated as shown in Equation 1:

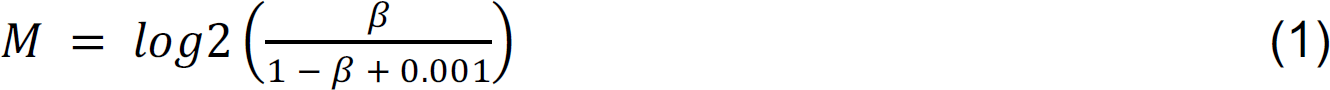

All statistical analysis was done in R. M-values of GATC sites within different genomic features were compared using the Kruskal-Wallis rank sum test, and each feature was compared to the genome-wide median M-value using a one-sample Wilcoxon signed rank test with Benjamini-Hochberg correction for multiple comparisons^84^.

Ordination of samples was performed after normalizing ꞵ-values. Since ꞵ-values are dependent on coverage, samples with low coverage stand out when visualized with non-metric ordination methods. Therefore, before ordination, we normalized coverage and methylated read counts between samples to reduce the effects of coverage on ordination distance. First, the coverage of all samples was normalized using quantile normalization^85^. Then, a scaling equation was calculated for each sample using the simple linear regression line between coverage and normalized coverage. This scaling equation was then applied to each sample’s coverage and methylated read count, both of which were used to calculate normalized ꞵ-values. Ordination of samples was performed using the R package vegan v. 2.6-4^86^. Non-metric multidimensional scaling (NMDS) was used to visualize sample clustering based on a Euclidean distance matrix calculated from normalized ꞵ-values. PERMANOVA and PERMDISP2 were done with vegan functions adonis2 and betadisper, respectively.

Gene ontology enrichment analysis was performed using R package clusterProfiler v. 4.6.2 and the Bioconductor *E. coli* K-12 annotation v. 3.8.2^87–89^. Genes enriched for biological process ontologies were filtered by a q-value cutoff of 0.05, and redundant GO terms were removed using the clusterProfiler “simplify” method.

### Data and code availability

Newly generated sequencing data for the identification of nucleotide modifications can be downloaded from SRA, BioProject PRJNA912686. Code and data for reproducing this analysis are found at https://github.com/BehringerLab/Methylation. The CoMMA R package is available at https://github.com/carl-stone/CoMMA.

## Acknowledgments

We would like to thank W-C. Ho, H. Lee, M. Lynch, and K. Samerotte for their helpful comments. High-performance computing resources were provided by the Research Computing Core at Arizona State University and the Advanced Computing Center for Research and Education (ACCRE) at Vanderbilt University. This work was supported by Army Research Office grants W911NF-21-1-0161 (M.G.B.), National Institutes of Health grants F32GM123703 (M.G.B.), and start-up funding from Vanderbilt University and the Vanderbilt Center for Infection, Immunology, and Inflammation.

---

**Figure S1.**
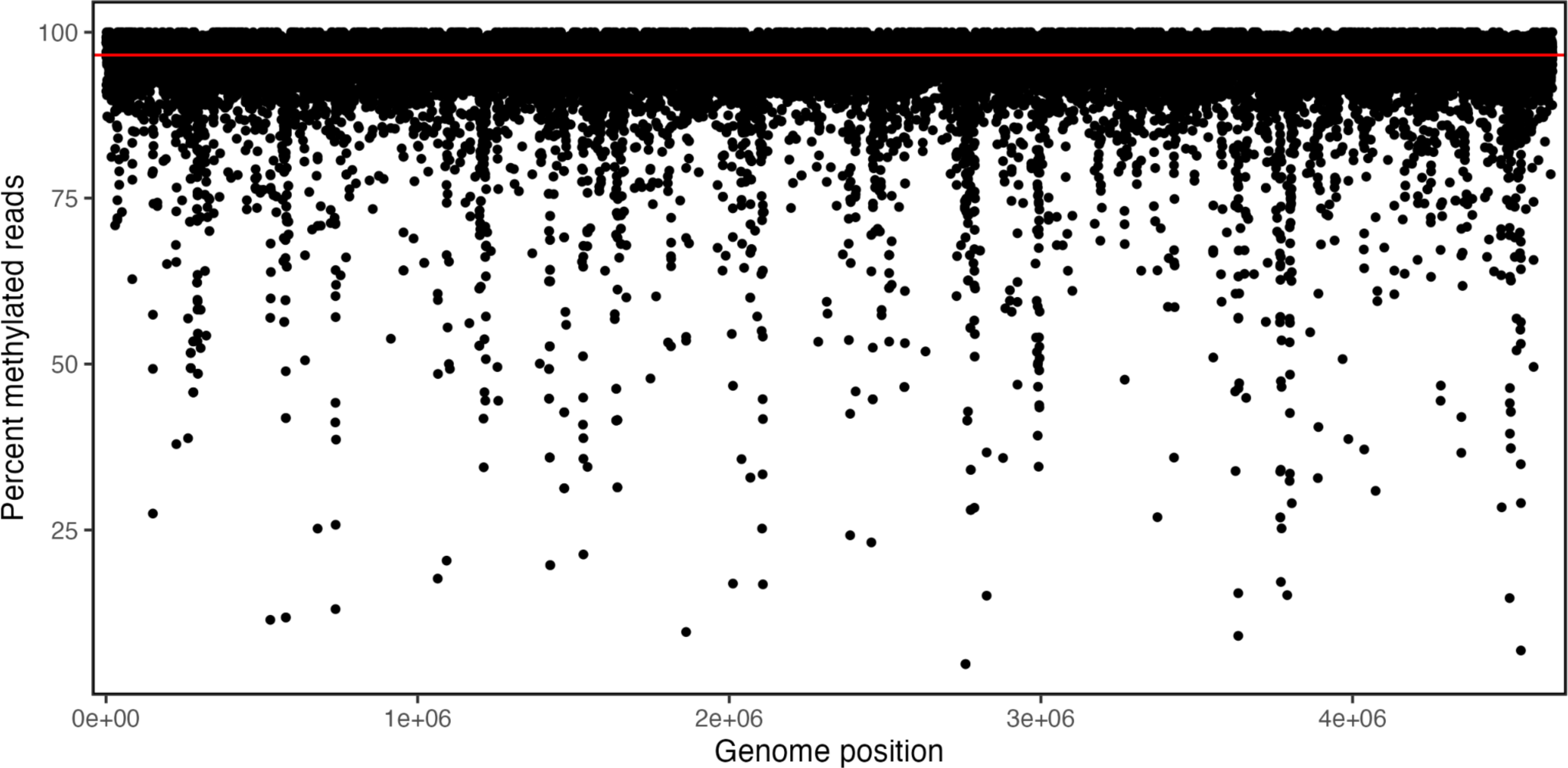
6mA methylation across the MG1655 genome. Each point represents the mean percent methylation of three sequencing replicates of the K-12 MG1655 ancestor strain at one GATC site. The genome-wide median methylation of 96.6% is shown for reference (red line).

**Figure S2.**
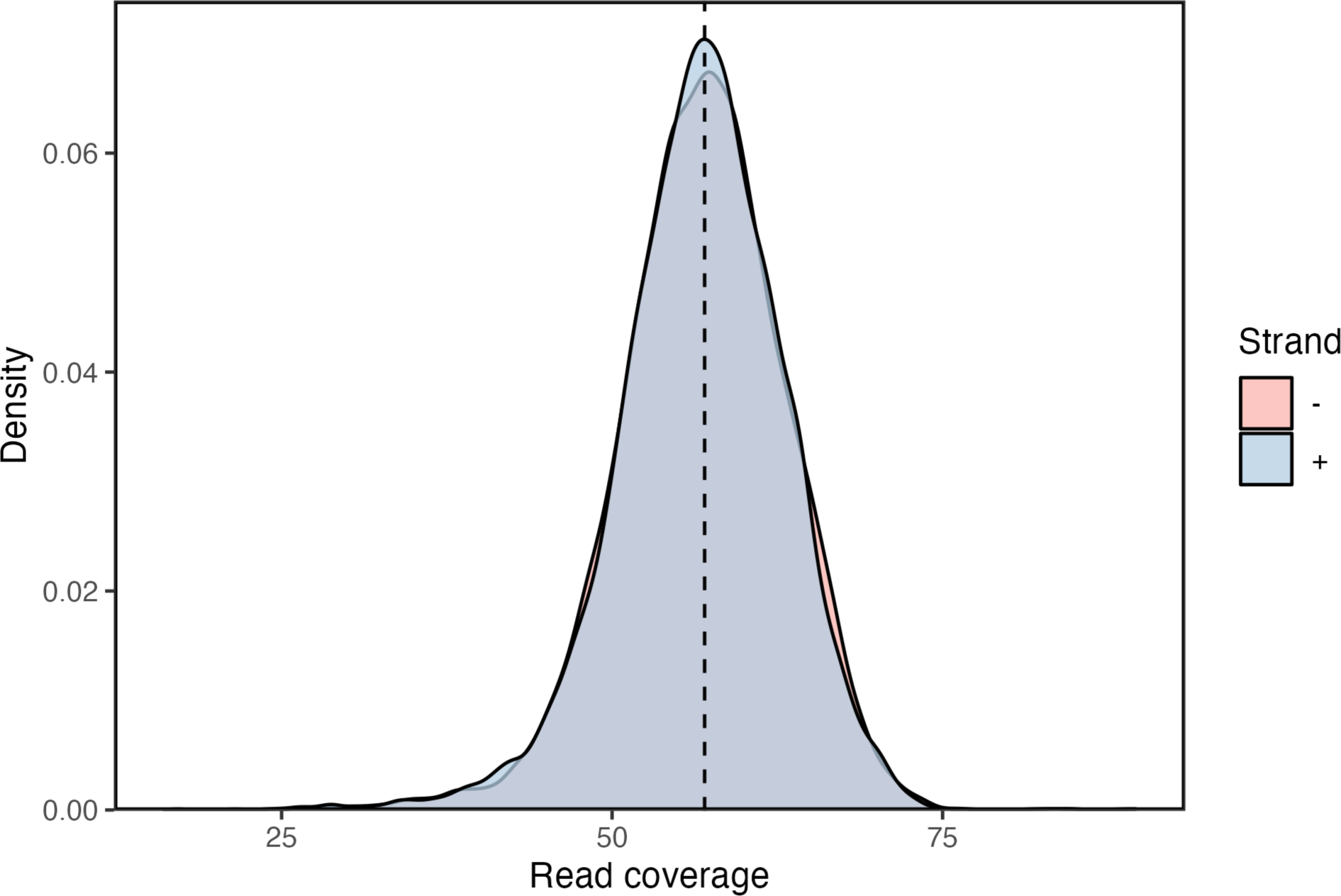
Sequencing coverage of GATC sites by Nanopore sequencing. Coverage distributions for each strand are shown separately, and both distributions have the same median of 57 reads (dashed line).

**Figure S3.**
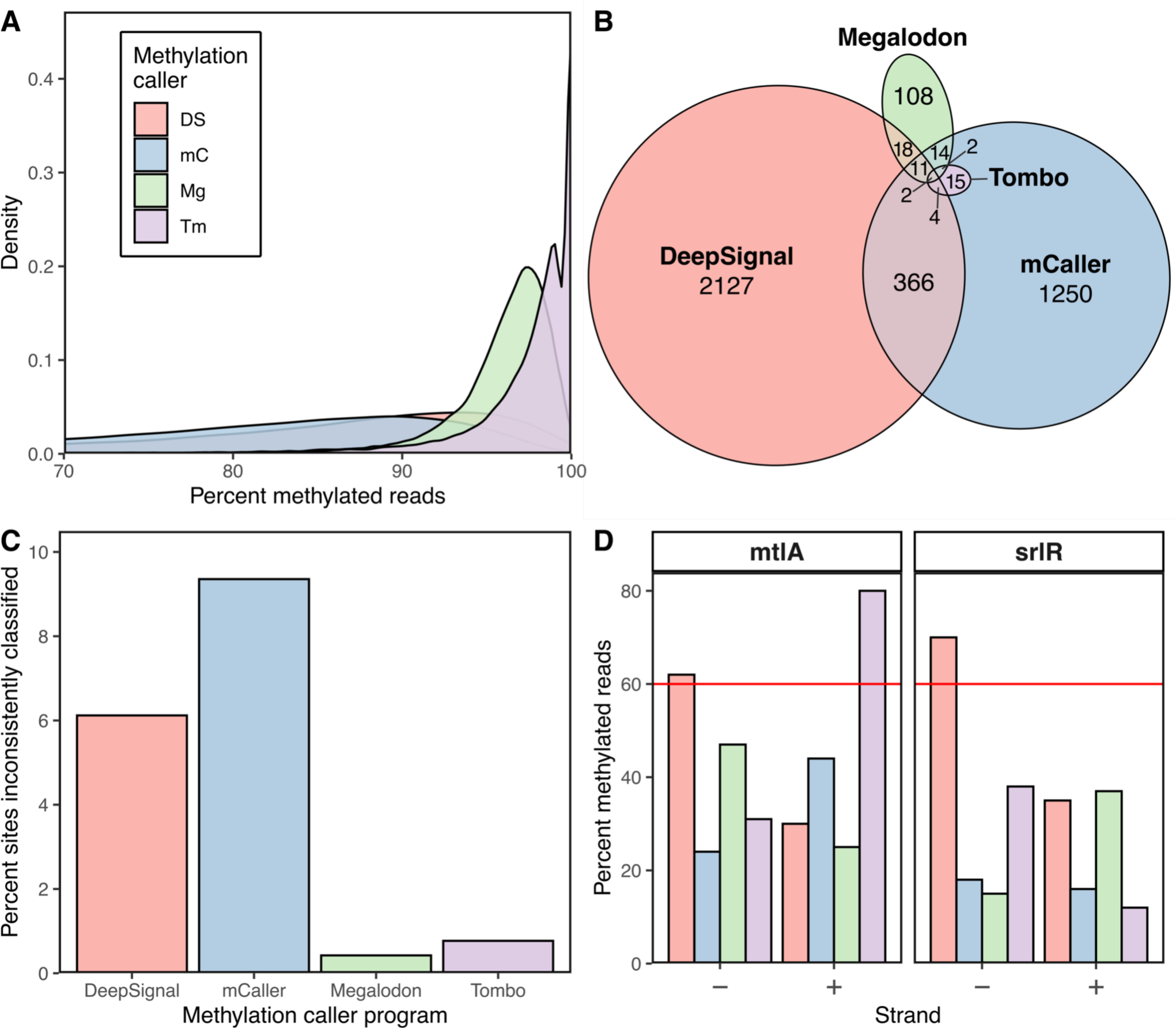
Comparison of bioinformatics programs on identifying methylated reads at GATC sites. (A) Distribution of percent of reads classified as methylated at all GATC sites in the genome for each caller. Dashed lines represent median percent methylation across all GATC sites for, from left to right, mCaller (83%), DeepSignal (87%), Megalodon (97%), and Tombo (99%). (B) Overlap between GATC sites classified as undermethylated by each program. Each area is labeled with the number of GATC sites. (C) Percentage of methylated GATC sites inconsistently classified as methylated across WT replicates for each program. (D) Percentage of reads classified as methylated by each program at GATC sites previously reported as <60% methylated.

**Figure S4.**
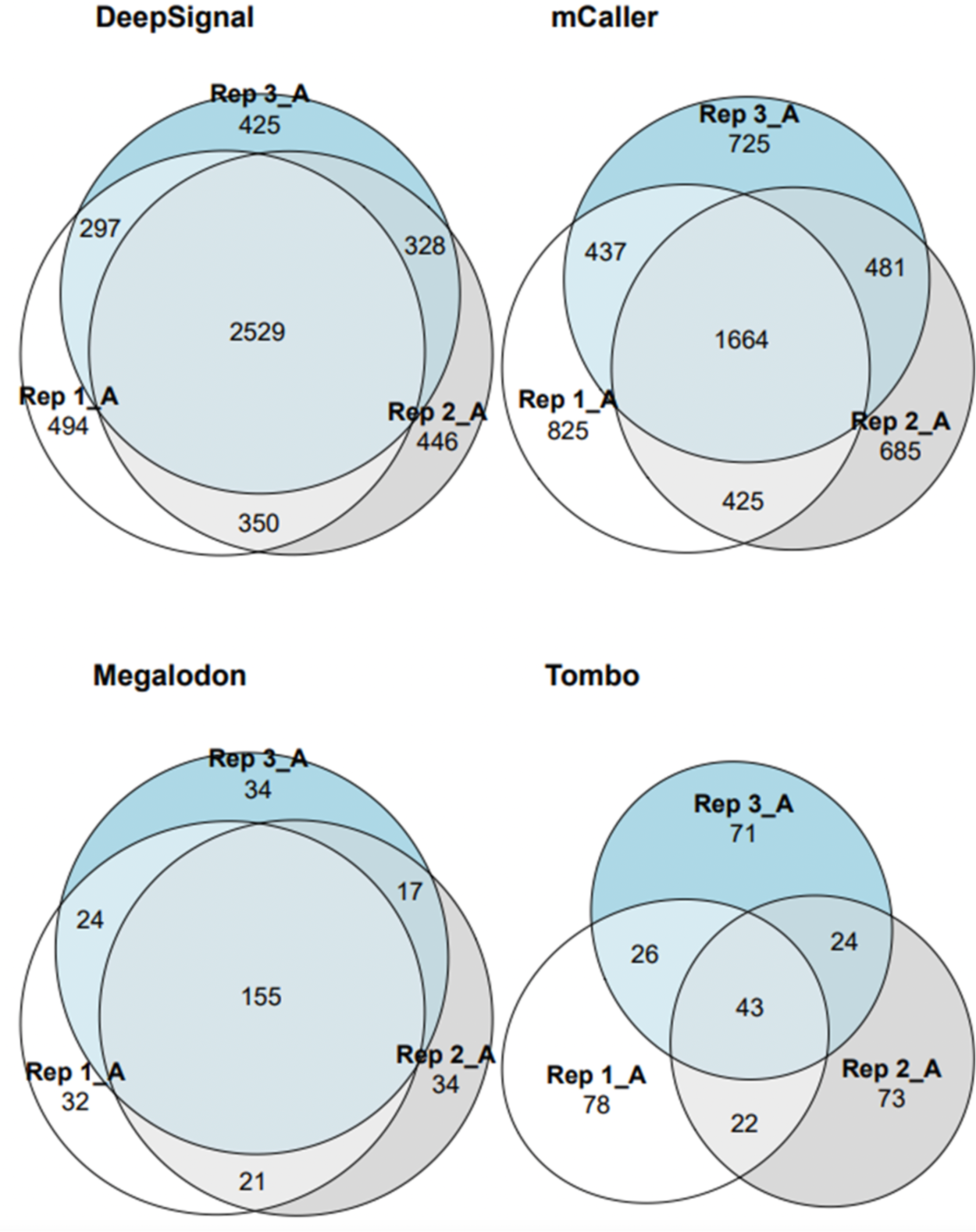
Overlap in sites identified as hypomethylated between three sequencing replicates for each caller. Each area is labeled with the number of GATC sites.

**Figure S5.**
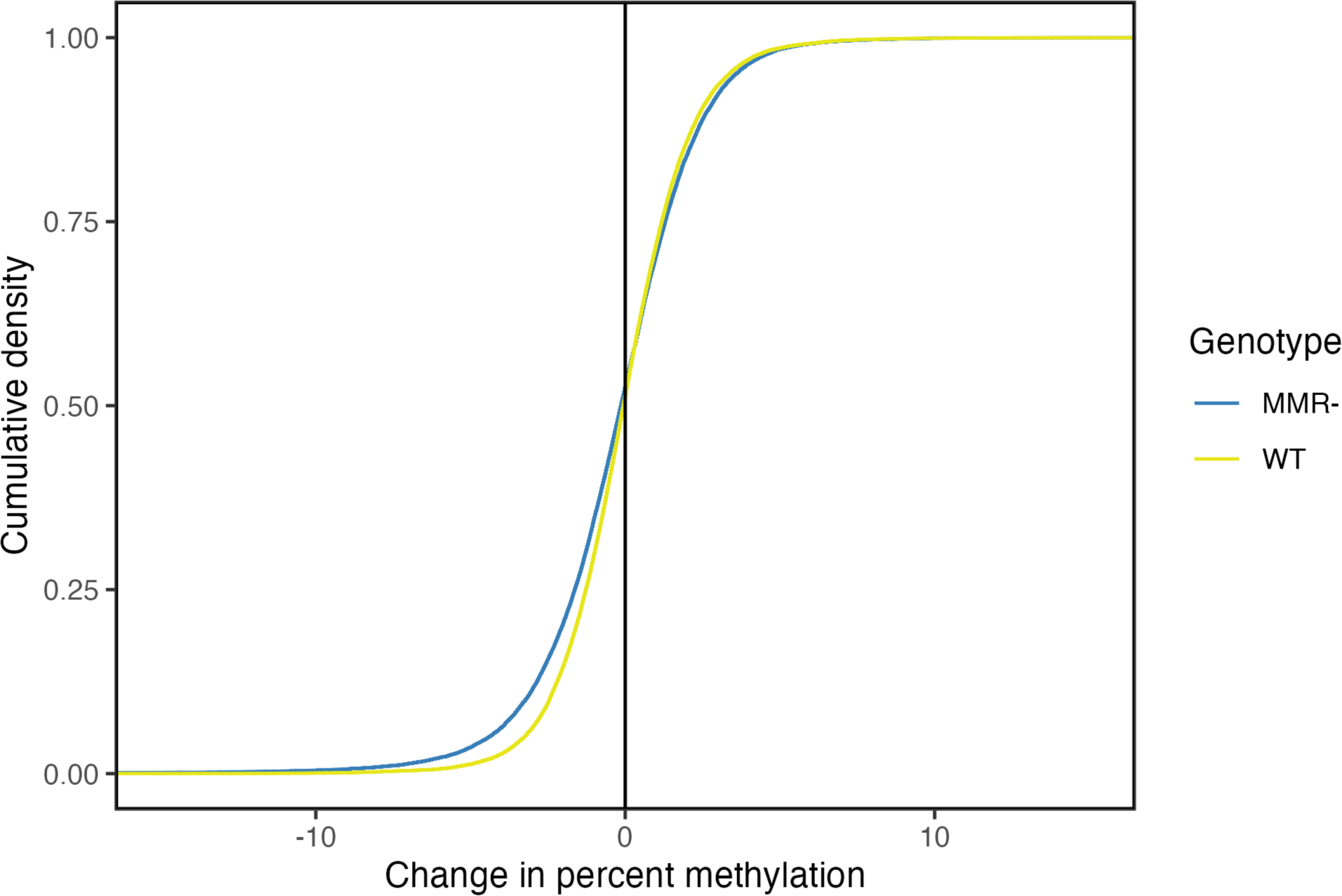
Distribution of changes in per-site methylation between evolved genotypes and their isogenic ancestor. For each GATC site, the difference between average percent methylation in the evolved genotype and in the ancestor was calculated, and values were used to calculate empirical cumulative density functions.

**Figure S6.**
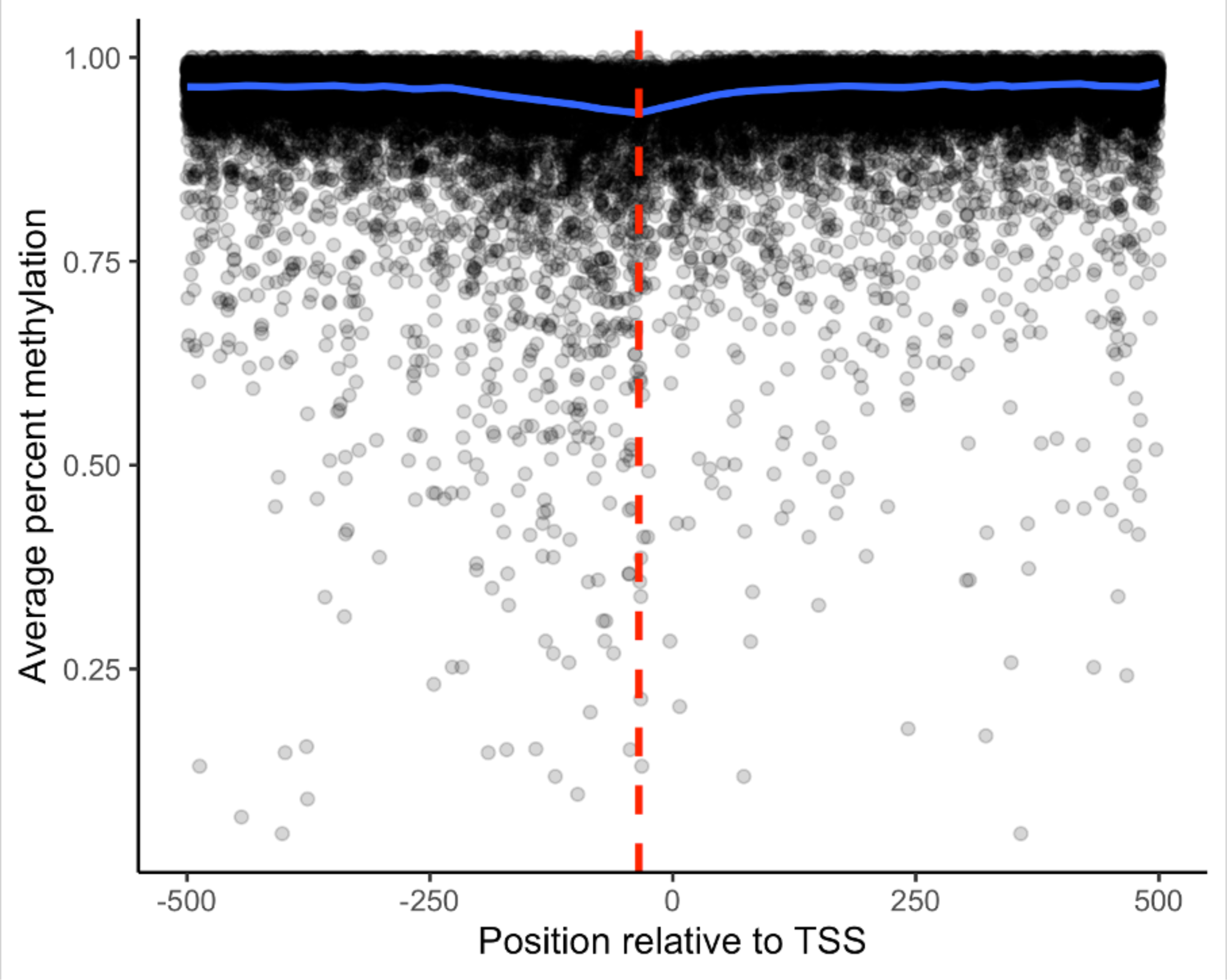
Methylation of all GATC sites relative to the transcription start site (TSS) Each point represents a single GATC site. The smoothed median was calculated by additive quantile regression smoothing (blue line), and the -35 position is shown for reference (dashed red line).

**Figure S7.**
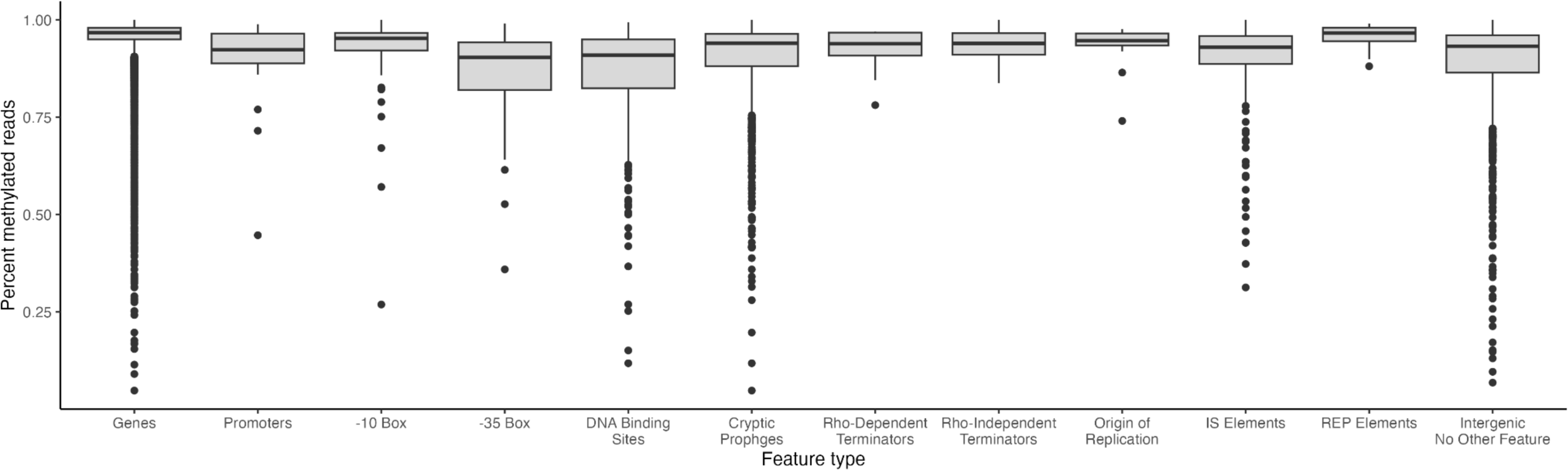
Average methylation at various genomic features. Boxplots show the median, first quartile, and third quartile of the percent methylated reads.

## REFERENCES

1. Sánchez-Romero, M. A. & Casadesús, J. The bacterial epigenome. Nat. Rev. Microbiol. 18, 7–20 (2020).

2. Wion, D. & Casadesús, J. N6-methyl-adenine: an epigenetic signal for DNA-protein interactions. Nat. Rev. Microbiol. 4, 183–192 (2006).

3. Seib, K. L., Srikhanta, Y. N., Atack, J. M. & Jennings, M. P. Epigenetic Regulation of Virulence and Immunoevasion by Phase-Variable Restriction-Modification Systems in Bacterial Pathogens. Annu. Rev. Microbiol. 74, 655–671 (2020).

4. Adhikari, S. & Curtis, P. D. DNA methyltransferases and epigenetic regulation in bacteria. FEMS Microbiol. Rev. 40, 575–591 (2016).

5. Ardissone, S. Cell Cycle Constraints and Environmental Control of Local DNA Hypomethylation in α-Proteobacteria. PLOS Genet (2016) doi:10.1371/journal.pgen.1006499.

6. Reisenauer, A., Kahng, L. S., McCollum, S. & Shapiro, L. Bacterial DNA Methylation: a Cell Cycle Regulator? J. Bacteriol. 181, 5135–5139 (1999).

7. Collier. A DNA methylation ratchet governs progression through a bacterial cell cycle. Proc Natl Acad Sci USA (2007) doi:10.1073/pnas.0708112104.

8. Sánchez-Romero, M. A., Cota, I. & Casadesús, J. DNA methylation in bacteria: from the methyl group to the methylome. Curr. Opin. Microbiol. 25, 9–16 (2015).

9. De Ste Croix, M., et al. Phase-variable methylation and epigenetic regulation by type I restriction-modification systems. FEMS Microbiol. Rev. 41, S3–S15 (2017).

10. Nye, T. M. et al. DNA methylation from a Type I restriction modification system influences gene expression and virulence in Streptococcus pyogenes. PLOS Pathog. 15, e1007841 (2019).

11. Oliveira, P. H. & Fang, G. Conserved DNA Methyltransferases: A Window into Fundamental Mechanisms of Epigenetic Regulation in Bacteria. Trends Microbiol. 29, 28–40 (2021).

12. Geier, G. E. & Modrich, P. Recognition sequence of the dam methylase of Escherichia coli K12 and mode of cleavage of Dpn I endonuclease. J. Biol. Chem. 254, 1408–1413 (1979).

13. Herman, G. E. & Modrich, P. Escherichia coli dam methylase. Physical and catalytic properties of the homogeneous enzyme. J. Biol. Chem. 257, 2605–2612 (1982).

14. Marinus, M. G. & Morris, N. R. Isolation of deoxyribonucleic acid methylase mutants of Escherichia coli K-12. J. Bacteriol. 114, 1143–1150 (1973).

15. Blow, M. J. et al. The Epigenomic Landscape of Prokaryotes. PLOS Genet. 12, e1005854 (2016).

16. Blattner, F. R. et al. The Complete Genome Sequence of *Escherichia coli* K-12. Science 277, 1453–1462 (1997).

17. Boye, E. & Løbner-Olesen, A. The role of dam methyltransferase in the control of DNA replication in E. coli. Cell 62, 981–989 (1990).

18. Pukkila, P. J., Peterson, J., Herman, G., Modrich, P. & Meselson, M. Effects of High Levels of DNA Adenine Methylation on Methyl-Directed Mismatch Repair in *Escherichia coli*. Genetics 104, 571– 582 (1983).

19. Schlagman, S. L., Hattman, S. & Marinus, M. G. Direct role of the Escherichia coli Dam DNA methyltransferase in methylation-directed mismatch repair. J. Bacteriol. 165, 896–900 (1986).

20. Messer, W. & Noyer-Weidner, M. Timing and targeting: The biological functions of Dam methylation in E. coli. Cell 54, 735–737 (1988).

21. Messer, W., Bellekes, U. & Lother, H. Effect of dam methylation on the activity of the E. coli replication origin, oriC. EMBO J. 4, 1327–1332 (1985).

22. Herman, G. E. Escherichia coli K-12 clones that overproduce dam methylase are hypermutable. J Bacteriol (1981) doi:10.1128/jb.145.1.644-646.1981.

23. Lahue, R. S. & Modrich, P. Methyl-directed DNA mismatch repair in Escherichia coli. Mutat. Res. Mol. Mech. Mutagen. 198, 37–43 (1988).

24. Roberts, D., Hoopes, B. C., McClure, W. R. & Kleckner, N. IS10 transposition is regulated by DNA adenine methylation. Cell 43, 117–130 (1985).

25. Tomcsanyi, T. & Berg, D. E. Transposition effect of adenine (Dam) methylation on activity of O end mutants of IS50. J. Mol. Biol. 209, 191–193 (1989).

26. Torreblanca, J. & Casadesús, J. DNA adenine methylase mutants of Salmonella typhimurium and a novel dam-regulated locus. Genetics 144, 15–26 (1996).

27. Heithoff, D. M., Sinsheimer, R. L., Low, D. A. & Mahan, M. J. An essential role for DNA adenine methylation in bacterial virulence. Science 284, 967–970 (1999).

28. Lu. SeqA: A negative modulator of replication initiation in E. coli. Cell (1994) doi:10.1016/0092-8674(94)90156-2.

29. Blyn, L. B., Braaten, B. A. & Low, D. A. Regulation of pap pilin phase variation by a mechanism involving differential dam methylation states. EMBO J. 9, 4045–4054 (1990).

30. Jakomin, M., Chessa, D., Bäumler, A. J. & Casadesús, J. Regulation of the Salmonella enterica std Fimbrial Operon by DNA Adenine Methylation, SeqA, and HdfR. J. Bacteriol. 190, 7406–7413 (2008).

31. Hernday, A. D., Braaten, B. A. & Low, D. A. The Mechanism by which DNA Adenine Methylase and PapI Activate the Pap Epigenetic Switch. Mol. Cell 12, 947–957 (2003).

32. van der Woude, M. W. Phase variation: how to create and coordinate population diversity. Curr. Opin. Microbiol. 14, 205–211 (2011).

33. Sánchez-Romero, M. A. & Casadesús, J. Waddington’s Landscapes in the Bacterial World. Front. Microbiol. 12, 685080 (2021).

34. Tavazoie, S. & Church, G. M. Quantitative whole-genome analysis of DNA-protein interactions by in vivo methylase protection in E. coli. Nat. Biotechnol. 16, 566–571 (1998).

35. Li, G.-M. Mechanisms and functions of DNA mismatch repair. Cell Res. 18, 85–98 (2008).

36. Lu, A. L., Clark, S. & Modrich, P. Methyl-directed repair of DNA base-pair mismatches in vitro. Proc. Natl. Acad. Sci. U. S. A. 80, 4639–4643 (1983).

37. Lee, H., Popodi, E., Tang, H. & Foster, P. L. Rate and molecular spectrum of spontaneous mutations in the bacterium Escherichia coli as determined by whole-genome sequencing. Proc. Natl. Acad. Sci. 109, E2774–E2783 (2012).

38. Posnick, L. M. & Samson, L. D. Influence of S-Adenosylmethionine Pool Size on Spontaneous Mutation, Dam Methylation, and Cell Growth of Escherichia coli. J. Bacteriol. 181, 6756–6762 (1999).

39. Westphal, L. L., Sauvey, P., Champion, M. M., Ehrenreich, I. M. & Finkel, S. E. Genomewide Dam Methylation in Escherichia coli during Long-Term Stationary Phase. mSystems 1, e00130–16 (2016).

40. Walworth, N. G. et al. Long-Term m5C Methylome Dynamics Parallel Phenotypic Adaptation in the Cyanobacterium Trichodesmium. Mol. Biol. Evol. 38, 927–939 (2021).

41. Oxford Nanopore Technologies. Megalodon. (2022).

42. Yuen, Z. W.-S. et al. Systematic benchmarking of tools for CpG methylation detection from nanopore sequencing. Nat. Commun. 12, 3438 (2021).

43. Liu. Detection of DNA base modifications by deep recurrent neural network on Oxford Nanopore sequencing data. Nat Commun (2019) doi:null.

44. Ni, P. et al. DeepSignal: detecting DNA methylation state from Nanopore sequencing reads using deep-learning. Bioinformatics 35, 4586–4595 (2019).

45. Oxford Nanopore Technologies. Tombo.

46. Wang, M. X. & Church, G. M. A whole genome approach to in vivo DNA-protein interactions in E. coli. Nature 360, 606–610 (1992).

47. van der Woude, M., Hale, W. B. & Low, D. A. Formation of DNA Methylation Patterns: Nonmethylated GATC Sequences in gut and papOperons. J. Bacteriol. 180, 5913–5920 (1998).

48. Tierrafría, V. H. et al. RegulonDB 11.0: Comprehensive high-throughput datasets on transcriptional regulation in Escherichia coli K-12. Microb. Genomics 8, (2022).

49. Cohen. A role for the bacterial GATC methylome in antibiotic stress survival. Nat Genet (2016) doi:10.1038/ng.3530.

50. Sanchez-Romero. Contribution of DNA adenine methylation to gene expression heterogeneity in Salmonella enterica. Nucleic Acids Res (2020) doi:10.1093/nar/gkaa730.

51. Wang, X. et al. Cryptic prophages help bacteria cope with adverse environments. Nat. Commun. 1, 147 (2010).

52. Ramisetty, B. C. M. & Sudhakari, P. A. Bacterial ‘Grounded’ Prophages: Hotspots for Genetic Renovation and Innovation. Front. Genet. 10, (2019).

53. Siguier, P., Gourbeyre, E. & Chandler, M. Bacterial insertion sequences: their genomic impact and diversity. FEMS Microbiol. Rev. 38, 865–891 (2014).

54. Lee, H., Doak, T. G., Popodi, E., Foster, P. L. & Tang, H. Insertion sequence-caused large-scale rearrangements in the genome of *Escherichia coli*. Nucleic Acids Res. gkw647 (2016) doi:10.1093/nar/gkw647.

55. Naas, T. Insertion sequence-related genetic variation in resting Escherichia coli K-12. Genetics (1994) doi:10.1093/genetics/136.3.721.

56. Behringer, M. G., et al. Escherichia coli cultures maintain stable subpopulation structure during long-term evolution. Proc. Natl. Acad. Sci. 115, E4642–E4650 (2018).

57. Akalin, A. et al. methylKit: a comprehensive R package for the analysis of genome-wide DNA methylation profiles. Genome Biol. 13, R87 (2012).

58. Qiao, J. et al. Construction of an Escherichia coli Strain Lacking Fimbriae by Deleting 64 Genes and Its Application for Efficient Production of Poly(3-Hydroxybutyrate) and l-Threonine. Appl. Environ. Microbiol. 87, e00381–21 (2021).

59. Henderson, I. R., Meehan, M. & Owen, P. A novel regulatory mechanism for a novel phase-variable outer membrane protein of Escherichia coli. Adv. Exp. Med. Biol. 412, 349–355 (1997).

60. Graveline. Lrp-DNA complex stability determines the level of ON cells in type P fimbriae phase variation. Mol Microbiol (2011) doi:10.1111/j.1365-2958.2011.07761.x.

61. Wallecha. Dam- and OxyR-dependent phase variation of agn43: essential elements and evidence for a new role of DNA methylation. J Bacteriol (2002) doi:10.1128/jb.184.12.3338-3347.2002.

62. Stevenson, G. et al. Structure of the O antigen of Escherichia coli K-12 and the sequence of its rfb gene cluster. J. Bacteriol. 176, 4144–4156 (1994).

63. Eguchi, Y. & Utsumi, R. Alkali Metals in Addition to Acidic pH Activate the EvgS Histidine Kinase Sensor in Escherichia coli. J. Bacteriol. 196, 3140–3149 (2014).

64. Kern, R., Malki, A., Abdallah, J., Tagourti, J. & Richarme, G. Escherichia coli HdeB Is an Acid Stress Chaperone. J. Bacteriol. 189, 603–610 (2007).

65. Yasui, R., Washizaki, A., Furihata, Y., Yonesaki, T. & Otsuka, Y. AbpA and AbpB provide anti-phage activity in *Escherichia coli*. Genes Genet. Syst. 89, 51–60 (2014).

66. Horton, J. R., Liebert, K., Bekes, M., Jeltsch, A. & Cheng, X. Structure and Substrate Recognition of the Escherichia coli DNA Adenine Methyltransferase. J. Mol. Biol. 358, 559–570 (2006).

67. Maier, J. A. H. & Jeltsch, A. Design and Application of 6mA-Specific Zinc-Finger Proteins for the Readout of DNA Methylation. in Zinc Finger Proteins: Methods and Protocols (ed. Liu, J.) 29–41 (Springer, 2018). doi:10.1007/978-1-4939-8799-3_3.

68. Sera, T. & Uranga, C. Rational design of artificial zinc-finger proteins using a nondegenerate recognition code table. Biochemistry 41, 7074–7081 (2002).

69. Nabel, C. S., Manning, S. A. & Kohli, R. M. The Curious Chemical Biology of Cytosine: Deamination, Methylation,and Oxidation as Modulators of Genomic Potential. ACS Chem. Biol. 7, 20–30 (2012).

70. Shapiro, R. & Klein, R. S. The Deamination of Cytidine and Cytosine by Acidic Buffer Solutions. Mutagenic Implications*. Biochemistry 5, 2358–2362 (1966).

71. Lindahl, T. & Nyberg, B. Heat-induced deamination of cytosine residues in deoxyribonucleic acid. Biochemistry 13, 3405–3410 (1974).

72. O’Brown, Z. K. & Greer, E. L. N6-Methyladenine: A Conserved and Dynamic DNA Mark. in DNA Methyltransferases - Role and Function (eds. Jeltsch, A. & Jurkowska, R. Z.) 213–246 (Springer International Publishing, 2016). doi:10.1007/978-3-319-43624-1_10.

73. Basilio, C., Wahba, A. J., Lengyel, P., Speyer, J. F. & Ochoa, S. Synthetic polynucleotides and the amino acid code, V*. Proc. Natl. Acad. Sci. 48, 613–616 (1962).

74. Sater, M. R. A. et al. DNA Methylation Assessed by SMRT Sequencing Is Linked to Mutations in Neisseria meningitidis Isolates. PLOS ONE 10, e0144612 (2015).

75. Brunet. Fur-dam regulatory interplay at an internal promoter of the enteroaggregative Escherichia coli Type VI secretion sci1 gene cluster. J Bacteriol (2020) doi:10.1128/JB.00075-20.

76. Mehling. A Dam methylation mutant of Klebsiella pneumoniae is partially attenuated. FEMS Microbiol Lett. (2007) doi:10.1111/j.1574-6968.2006.00581.x.

77. McIntyre, A. B. R. et al. Single-molecule sequencing detection of N6-methyladenine in microbial reference materials. Nat. Commun. 10, 579 (2019).

78. Oxford Nanopore Technologies. Tombo Documentation Modified Base Detection. https://nanoporetech.github.io/tombo/modified_base_detection.html.

79. Cingolani, P. et al. A program for annotating and predicting the effects of single nucleotide polymorphisms, SnpEff. Fly (Austin) 6, 80–92 (2012).

80. Danecek, P. et al. The variant call format and VCFtools. Bioinformatics 27, 2156–2158 (2011).

81. Santos-Zavaleta, A. et al. RegulonDB v 10.5: tackling challenges to unify classic and high throughput knowledge of gene regulation in E. coli K-12. Nucleic Acids Res. 47, D212–D220 (2019).

82. Wang, H.-Q., Tuominen, L. K. & Tsai, C.-J. SLIM: a sliding linear model for estimating the proportion of true null hypotheses in datasets with dependence structures. Bioinforma. Oxf. Engl. 27, 225–231 (2011).

83. Du, P. et al. Comparison of Beta-value and M-value methods for quantifying methylation levels by microarray analysis. BMC Bioinformatics 11, 587 (2010).

84. Benjamini, Y. & Hochberg, Y. Controlling the False Discovery Rate: A Practical and Powerful Approach to Multiple Testing. J. R. Stat. Soc. Ser. B Methodol. 57, 289–300 (1995).

85. Bolstad, B. M., Irizarry, R. A., Åstrand, M. & Speed, T. P. A comparison of normalization methods for high density oligonucleotide array data based on variance and bias. Bioinformatics 19, 185–193 (2003).

86. Oksanen, J., et al. vegan: Community Ecology Package. (2022).

87. Wu, T. et al. clusterProfiler 4.0: A universal enrichment tool for interpreting omics data. The Innovation 2, 100141 (2021).

88. Yu, G., Wang, L.-G., Han, Y. & He, Q.-Y. clusterProfiler: an R Package for Comparing Biological Themes Among Gene Clusters. OMICS J. Integr. Biol. 16, 284–287 (2012).

89. Carlson, M. org.EcK12.eg.db: Genome wide annotation for E coli strain K12. (2019).

