## Supplemental Text 1 for "Differential adenine methylation analysis reveals increased variability in 6mA in the absence of methyl-directed mismatch repair"

**Supplementary Text 1**

**Evaluation of methylation callers revealed large differences in precision.**

Methylation-calling programs for Nanopore data have been thoroughly benchmarked on the classification of 5mC. While popular programs can identify 6mA, there are limited studies where Nanopore data has been used on 6mA, especially in bacteria. Thus, we identified the location of the 38,240 GATC sites in the E. coli K-12 substr. MG1655 genome (**Fig S1**) and compared the reliability and reproducibility of commonly used methylation callers in characterizing 6mA at these sites. We sequenced DNA from three equal aliquots of a single *E. coli* K-12 MG1655 (strain PMF2 from Lee *et al.* 2012) overnight culture on an ONT MinION portable sequencer[1]. Nanopore sequencing produced an average read depth of ~50 reads per GATC site, and there was no significant strand bias in coverage (**Fig. S2**). We then used four different software packages (DeepSignal, mCaller, Megalodon, and Tombo) to calculate the proportion of methylated reads to total reads, called the percent methylation for each GATC site (**Dataset S1**)[2–6]. All four of these packages differ in their approaches and dependencies.

Each methylation caller produced varying distributions of predicted methylation values across all GATC sites (**Fig. S3A)**. Tombo classified a high proportion of sites as completely methylated (all reads were classified as methylated at 31.8% of sites). This is consistent with previous benchmarking on 5mC methylation where Tombo over-predicted methylation at CpG sites[7]. DeepSignal and mCaller predicted fewer sites as methylated (median percent methylated reads of 87% and 83%, respectively). Megalodon identified a median percent methylation of 97% across all sites.

We next determined the precision of each methylation caller by comparing how consistently each GATC site was called as methylated across the three sequencing replicates. We considered a GATC site to be completely methylated if ≥60% of reads on both strands were inferred to be methylated. Alternatively, GATC sites with fewer than 60% of reads methylated on one or both strands were classified as hemimethylated or hypomethylated, respectively. Each caller identified different numbers of hypomethylated sites, and there is little overlap between the specific sites classified as hypomethylated by different callers (**Fig. S3B**). Across the 38,240 single-stranded GATC sites in the *E. coli* K-12 substr. MG1655 genome, Megalodon was the most precise, producing the fewest number of inconsistently classified GATC sites (162 sites, or 0.4%) outperforming Tombo, the second most consistent methylation caller with 294 inconsistent sites (**Fig. S3C; Table S1**). DeepSignal and mCaller were the most inconsistent classifiers with 2340 and 3577 sites, respectively, inconsistently identified between sequencing replicates.

Given the high activity of Dam methyltransferase and the central role of GmATC in directing mismatch repair, GATC sites in *E. coli* are expected to be fully methylated except for a few key loci[8,9]. Thus, we compared how well each caller identified known undermethylated sites as a measure of caller accuracy. Previous estimates that measured 6mA by cutting with methylation-sensitive and -insensitive endonucleases have suggested that about 0.1% of *E. coli* GATC sites (or ~20 double-stranded GATC sites) are consistently found in a completely hypomethylated state *in vivo*[8,10]. In our analysis, Megalodon identified 22 consistently hypomethylated GATC sites across the three sequencing replicates. Tombo called 1 site as hypomethylated, while DeepSignal and mCaller identified 78 and 41 sites, respectively (**Table S1**). Based on these results, Megalodon is the most accurate as it identifies methylated sites at similar frequencies to those seen in previously published work[8].

As another measure of accuracy, we next examined how each caller identifies known undermethylated sites in the genome by focusing on the *srl* and *mtl* operons which are necessary for the import and metabolism of sorbitol and mannitol, respectively. Previous studies have established that in the absence of sorbitol and mannitol, transcriptional repressors bind to regions in the promoters that overlap with GATC motifs, preventing Dam methyltransferase binding and resulting in consistent undermethylation at these GATC sites (**Table S2**)[8,9,11]. Of the four methylation callers, only mCaller and Megalodon correctly identified both sites as hypomethylated (**Fig. S3D**). Tombo identified the *srl* site as hypomethylated and the *mtl* site as hemimethylated, while DeepSignal called both sites as hemimethylated. Taken together, these data show that Megalodon is the most specific and accurate caller when classifying the methylation status of GATC sites using Nanopore data.

### References


1. Lee, H., Popodi, E., Tang, H. & Foster, P. L. Rate and molecular spectrum of spontaneous mutations in the bacterium Escherichia coli as determined by whole-genome sequencing. *Proc. Natl. Acad. Sci.* **109**, E2774–E2783 (2012).
2. Ni, P. *et al.* DeepSignal: detecting DNA methylation state from Nanopore sequencing reads using deep-learning. *Bioinformatics* **35**, 4586–4595 (2019).
3. Oxford Nanopore Technologies. Tombo. https://github.com/nanoporetech/tombo.
4. Oxford Nanopore Technologies. Tombo Documentation Modified Base Detection. https://nanoporetech.github.io/tombo/modified_base_detection.html.
5. Oxford Nanopore Technologies. Megalodon. (2022).
6. McIntyre. Single-molecule sequencing detection of N6-methyladenine in microbial reference materials. *Nat Commun* (2019) doi:10.1038/s41467-019-08289-9.
7. Yuen, Z. W.-S. *et al.* Systematic benchmarking of tools for CpG methylation detection from nanopore sequencing. *Nat. Commun.* **12**, 3438 (2021).
8. Tavazoie, S. & Church, G. M. Quantitative whole-genome analysis of DNA-protein interactions by in vivo methylase protection in E. coli. *Nat. Biotechnol.* **16**, 566–571 (1998).
9. Cohen, N. R. *et al.* A role for the bacterial GATC methylome in antibiotic stress survival. *Nat. Genet.* **48**, 581–586 (2016).
10. Wang, M. X. & Church, G. M. A whole genome approach to in vivo DNA-protein interactions in E. coli. *Nature* **360**, 606–610 (1992).
11. van der Woude, M., Hale, W. B. & Low, D. A. Formation of DNA Methylation Patterns: Nonmethylated GATC Sequences in gut and papOperons. *J. Bacteriol.* **180**, 5913–5920 (1998).
